# A hemispheric dome setup for naturalistic visual stimulation in head-fixed mice

**DOI:** 10.1101/2024.05.31.596599

**Authors:** Magdalena Kautzky, Verena Peterreins, Yongrong Qiu, Zhijian Zhao, Ann H. Kotkat, Steffen Katzner, Thomas Euler, Laura Busse

## Abstract

The visual system of any animal species is adapted to its ecological niche. Thus, investigating visual function and behavior using naturalistic stimuli holds significant potential. In mice, these adaptations include a field of view of ∼280° and cone opsins sensitive to UV and green wavelengths. Such adaptations, however, cannot be probed with standard consumer displays. To present naturalistic visual stimuli to mice, we built a hemispheric dome setup, enabling the controlled projection of wide-field movies with UV-green spectral content. For our UV-green projection, we used a customized light engine with external high-power LEDs. We mitigated spatial distortions introduced by the projection through a geometry-independent calibration procedure. Additionally, we adapted a head-mounted eye tracking system to capture behavioral responses of head-fixed mice viewing these stimuli. We validated our setup by quantifying the pupillary light reflex to uniform stimuli and the optokinetic reflex to drifting gratings. Finally, in experiments with naturalistic movies, we investigated whether mice showed differential saccades and eye positions based on visual input. Comparing naturalistic movies to a uniform screen control condition, we observed that although head-fixed mice did not make targeted saccades during movies, their overall eye position consistently shifted towards more frontal regions of visual space. This indicates that mice adjust their eye position in a stimulus-dependent way, potentially to optimize visual processing of information ahead in the visual field. Together, our results highlight the utility of our setup for in vivo studies of the mouse visual system with more naturalistic visual stimulation.

**Significance Statement:** Over the past decades, mice have emerged as a prominent model for vision. Nonetheless, compared to primates, mice inhabit a different visual niche, featuring several species-specific adaptations, including a wide field of view and sensitivity to UV light. Both adaptations cannot be adequately studied using consumer displays commonly employed in vision research. We therefore built a hemispheric dome setup allowing the presentation of wide field movies with appropriate spectral content and tracking eye movements and pupil dynamics in head-fixed mice with minimal shadows. We provide a baseline assessment of the setup leveraging two well-established eye reflexes. We also showcase the setup’s utility by reporting adjustments in eye movements when mice viewed the naturalistic movies.

## Introduction

The visual systems of animals are adapted by evolutionary pressures to the statistics of their natural habitats and behavioral needs (Baden et al., 2020; Land, 2015; Sedigh-Sarvestani and Fitzpatrick, 2022). Notably, neuronal populations in the visual system demonstrate preferences aligned with the statistics of natural scenes (reviewed in Geisler, 2007; Sedigh-Sarvestani and Fitzpatrick, 2022). One of the most-studied examples is the dominance of cardinal orientations in natural scenes (Switkes et al., 1978; van der Schaaf and van Hateren, 1996; Coppola et al., 1998; Girshick et al., 2011; Betsch et al., 2004), which is mirrored by a cardinal bias in neuronal orientation preference (e.g., Yacoub et al., 2008; Nasr and Tootell, 2012; Shen et al., 2014; Li et al., 2003; Coppola et al., 1998; Kreile et al., 2011).

Mice exhibit distinct adaptations in their visual system, notably a wide field of view (FOV) and the retinal gradient for UV and green sensitivity. With laterally placed eyes providing a panoramic view of ∼ 280 ° (Seabrook et al., 2017; Pinke et al., 2023), the mouse senses the visual information from its surroundings by its rod and two types of cone photoreceptors (Haverkamp, 2005). The cones express opsins sensitive to UV (S-opsin, peak 360 nm) and green (M-opsin, peak 510 nm) (Jacobs et al., 2004). The “true” S-cones (5% of the cones) exclusively express the S-opsin and are distributed throughout the retina, albeit with a “hotspot“ located in the ventral retina (Nadal-Nicolás et al., 2020). The M-cones (95%) importantly contribute to this ventral dominance of UV-sensitivity by co-expressing S-opsin, with higher co-expression ratios towards the ventral retina (Applebury et al., 2000; Röhlich et al., 1994; Haverkamp, 2005; Baden et al., 2013). This dorso-ventral UV-green gradient reflects the visual environment statistics for mice (Qiu et al., 2021; Abballe and Asari, 2022), potentially aiding in the detection of dark contrasts in the sky region (Qiu et al., 2021; Baden et al., 2013).

A promising approach for probing the function of the visual system is to use naturalistic, species-specific stimuli. Such stimuli, however, have so far rarely been used for the mouse, given limitations in both the visual stimulus material and its presentation. While several studies, inspired by previous work in cats (Betsch et al., 2004), have acquired footage from head-mounted cameras in mice roaming in a cage or arena (e.g., Froudarakis et al., 2014; Hofer et al., 2011), these movies were only gray-scale and did not consider the effective image-stabilization mechanisms through eye movements (Meyer et al., 2018, 2020). Also, visual stimuli for mice are often presented on small off-the-shelf RGB consumer displays, which cannot activate the S-opsin and typically cover only a small fraction of the mouse visual field.

Recently, some of the technical limitations regarding stimulus material and presentation have been overcome. Two data sets with movies or images of outdoor mouse environments have been acquired with custom-made UV-/green sensitive cameras and a large FOV, mimicking some aspects of the mouse visual system (Qiu et al., 2021; Abballe and Asari, 2022). Furthermore, several solutions for visual stimulation in the UV spectrum have been proposed (Rhim et al., 2017; Franke et al., 2019; Denman et al., 2017; Tan et al., 2015), including an arbitrary-spectrum spatial visual stimulator (Franke et al., 2019). Finally, immersive visual stimulation, including from the overhead space, can now be provided in hemispheric dome setups (Shapcott et al., 2022; Lopes et al., 2021). So far, however, these elements have not yet been combined to present a naturalistic visual environment for mice.

Here, we implemented a hemispheric dome setup to present naturalistic visual stimuli with the appropriate spectral content to head-fixed mice and measure their behavioral responses. We describe the components of our setup and provide robust methods for spatial calibration. We validated that we achieved sufficient intensity of stimulation by measuring the pupillary light reflex (PLR). Using the optokinetic reflex (OKR), we verified that our methods for measuring and quantifying eye movements in our setup have appropriate precision. Exploiting the naturalistic movies from our previous work (Qiu et al., 2021), we confirmed that head-fixed mice do not use saccadic eye movements to orient to specific features of the scenes. Interestingly, however, we found that mice systematically shifted eye position to more frontal regions of the visual field when viewing naturalistic movies compared to a mean-luminance gray screen. This finding adds to the growing evidence that mice perform complex visuo-motor adjustments according to visual input, potentially to maximize coverage of the visual field just ahead.

## Materials and Methods

### Animals

All procedures complied with the European Communities Council Directive 2010/63/EU and the German Law for Protection of Animals, and were approved by local authorities following appropriate ethics review (license number ROB-55.2-2532.Vet 02-18-190).

The experiments were conducted with six PVCre mice of either sex (B6;129P2-Pvalbtm1(cre)Arbr/J; #008069, Jackson Laboratory) of 3, 8, or 11 months of age at the beginning of the experiment. They were housed on a 12/12 h light-dark cycle, with experiments being conducted in their light phase.

### Surgical procedures

Thirty minutes before the start of surgery, mice were injected subcutaneously with an analgesic (Metamizole, 200 mg/kg, sc, MSD Animal Health, Brussels, Belgium). Isoflurane (5% in oxygen, CP-Pharma, Burgdorf, Germany) anesthesia was slowly induced in a chamber. After induction, mice were fixed in a stereotaxic frame (Drill and Microinjection Robot, Neurostar, Tübingen, Germany), with isoflurane being delivered through a face mask. The mice were placed onto a closed-loop heat plate (ATC 1000, WPI, Berlin, Germany), to ensure a stable body temperature of 37° Celsius. The flow rate and isoflurane concentration (0.5%–2% in oxygen) were adjusted to ensure a stable depth of anesthesia, as judged by the absence of the pedal reflex.

The eyes were covered with Bepanthen (Bayer, Leverkusen, Germany), and the ointment was re-applied several times during surgery. Additional analgesic (Buprenorphine, 0.1 mg/kg, sc, Bayer, Leverkusen, Germany) was administered. The head was thoroughly depilated (Avon Skin-so-Soft, facial, Avon, Germany). For disinfection of the surgical area on the mouse’s head, an iodine solution (Braunodivon, 10%, B. Braun, Melsungen, Germany) was applied. After administration of a local anesthetic (Lidocaine hydrochloride, bela-pharm, Vechta, Germany), a skin incision was made with a scalpel and an oval piece of skin was removed above the midline. Subsequently, the skull was cleaned from any tissue residues, the periosteum was removed, and the bone was roughened by a drop of H_2_O_2_ (3%, AppliChem, Darmstadt, Germany). To ensure a skull-flat implantation of the headbar, four reference points (bregma, lambda, and two points 2 mm to the left and right of the midline, respectively) were measured, and, if required, the head was re-positioned.

The exposed and cleaned area of the skull was covered with OptiBond FL primer and adhesive (Kerr dental, Rastatt, Germany) and then hardened with UV light. A lightweight aluminium headbar (24 × 4 mm, **Fig. 1b, left**) was positioned centrally anterior to the ears above the skull and attached with dental cement (Ivoclar Vivadent, Ellwangen, Germany). Next, an aluminum camera holder (**Fig. 1b, left**) was implanted, whose bottom and side surfaces were filed before implantation to roughen the material for better adhesion. To ensure long-term stability and reproducibility of the camera position, but also minimal weight, the camera holder was a non-symmetrical, pentagon shaped aluminum block (side lengths: 500 x 700 x 280 x 500 x 530 mm). A small hole with an M2 thread was positioned on the side of the block pointing towards the back of the mouse. This hole held a 3 mm plastic screw (Polyamide MXD6) with a pan shaped head (CSPPNR-RENY-M2-3, Misumi, Mexico).

**Figure 1:**
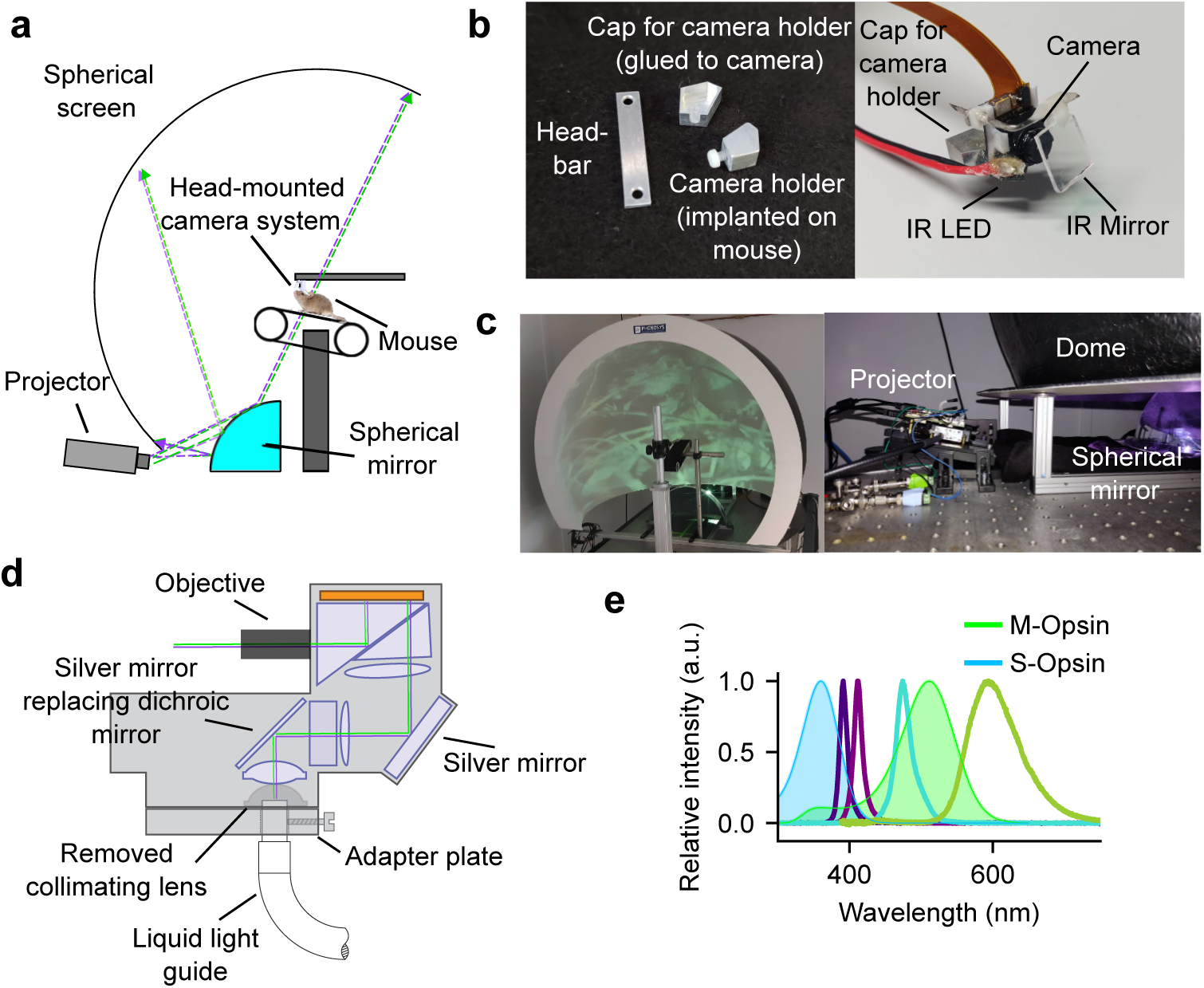
Hemispheric dome setup for visual stimulation and head-mounted camera system for eye tracking. (**a**) Schematic drawing of the dome setup with head-fixed mouse wearing the head-mounted camera system. (**b**) Implanted and removable parts of the eye tracking system. *Left*: The aluminum headbar next to the implanted pentagon-shaped aluminum camera holder (*bottom*) with a screw to clamp the cap holding the eye tracking system (*top*). *Right*: Front view of the head-mounted eye tracking system. (**c**) Dome and LC. *Left*: Dome setup with the LC projector, spherical mirror, and linear treadmill. *Right*: The LC connected to the liquid light guide and two output cables to trigger the UV and green LEDs of the light source and mounted on an angled platform. (**d**) Optical light path of the LC and custom modifications. (**e**) Spectra of LEDs used for UV and green stimulation from the 6-channel high intensity light source (Chrolis, Thorlabs; normalized intensities) and spectra of S- and M-opsin sensitivities of the mouse retina.

To correctly position the camera holder during implantation, a miniature eye tracking camera system (see below) was attached while the camera holder was lowered towards the skull. As soon as the position of the camera holder was appropriate to yield high-quality images of the eye, a thin layer of superglue was spread between headbar and camera holder, and the holder was further lowered onto the headbar. The surrounding spaces between camera holder and skull were filled with dental cement, which was also used to build a small rim around the sides of the camera holder.

At the end of the surgery, the edges of the wounds were treated with Braunodivon (10%, B. Braun, Melsungen, Germany). A long-term analgesic (Meloxicam, 2 mg/kg, sc, Boehringer Ingelheim, Ingelheim, Germany) was injected. For 1 mouse, to prevent infection, an antibiotic (Baytril, 5 mg/kg, sc, Bayer Animal Health GmbH, Leverkusen, Germany) was administered. The analgesic was given for 3 consecutive days and the antibiotics for 7 days post surgery. The health status of the mice was evaluated with a score sheet for at least 5 days after the surgery.

After the recovery period of one week, mice were accustomed to the setup and head fixation. During this time the focus and the mirror of the eye tracking camera (see below) were adjusted if necessary. After acclimatization, two recording sessions were carried out per day, for a maximum of one hour per session.

### Hemispheric dome setup, mirror and illumination

To present visual stimuli covering most of the large field of view of the mouse, we implemented a hemispheric dome setup, consisting of a screen, a mirror, and a projector, all fixed on a frame and a bread board table (**Fig. 1a, c**). The mouse was head-fixed in the center of the dome and was free to run on a linear treadmill (Speed belt, PhenoSys, Berlin, Germany).

The dimensions of the hemispheric screen (PhenoSys, Berlin, Germany) were 120 cm in width, 113 cm in depth and 106 cm in height. From the position of the mouse in the center of the dome, this corresponded to a width of 250° and a height of 160°, thus covering substantial portions of the visual field of mice. An acrylic spherical mirror (PhenoSys, Berlin, Germany), sized as a quarter sphere, illuminated the inner dome surface. The mirror was positioned on the table at the bottom of the hemispherical screen, allowing the projector to be fixed to the table behind and slightly below the dome. This setup prevented shadows of equipment and maintained a short distance between the optical components. The projector was centered onto the mirror, such that the center of the image was displayed in the center of the dome, directly in front of the mouse.

The projector was angled by about 13° (**Fig. 1c**) to sufficiently diverge the light hitting the outer areas of the mirror and thus ensure full illumination of the 250° wide dome.

To illuminate the dome, we used a Digital Light Processing (DLP) LightCrafter 4500 Fiber Couple projector (Fiber-E4500MKIITM, EKB Technologies Ltd., Israel; referred to as “LC”) (Franke et al., 2019). A 6-wavelength, high-power LED Source (Chrolis, Thorlabs, USA) was connected to the LC via a liquid light guide (LLG). For stimulation of the mouse S-opsin, two LEDs peaking at 385 nm and 405 nm were used, with a total maximum power of 2150 mW (measured at the end of the LLG; from now on referred to as “UV”). For stimulation of the M-opsin, a 475 nm LED and a 590 nm LED were combined, yielding a total light power of 770 mW (from now on referred to as “green”). To optimize the LC’s light transmission, we performed two custom modifications: (1) we removed the first collimating lens, and (2) we replaced the dichroic mirror by a silver mirror to improve reflection of all wavelengths, including the UV. Considering the output at the end of the LLG as 100%, we achieved transmission as measured at the LC’s front lens of 8% of light from the 385 nm LED, 14% of light from the 405 nm LED, and 33% of light from the 590 nm LED.

Stimuli were presented with 8 bit depth and at a resolution of 912×1440 pixels. To further increase stimulus intensity, the LC was configured to operate in “pattern mode”, which, amongst others, allows assigning each of the 24 bitplanes (3 RGB color channels × 8 bit) of every frame to an arbitrary combination of LEDs. Hence, restricting the video output to only two colors (green and UV), resulted in more time within a frame for the green and UV LEDs, and thus brighter images (Franke et al., 2019).

To relate the light intensity on dome surface to photoisomerization rates at the photoreceptor level, we used a power meter (**Table 1**, part #11) to measure the light power at the center of the dome for the different LEDs with different wavelengths (*λ*, in [nm]). We calculated the light power per unit area (*P_el_*, in [W/m^2^]), by dividing the measured light power by the sensor area *A*_stim_ = 10*^−^*^4^m^2^. We then converted light power to photon flux (*P_P hi_*, in [photons/s/m^2^]) using:

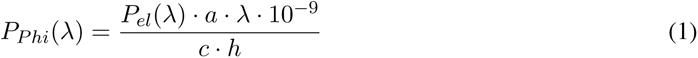

with constants *a* = 6.242 · 10^18^ *eV/J*, the speed of light *c* = 299, 792, 458 m/s, and Planck constant *h* = 4.135667 · 10^−15^ *eV* · s. We next estimated the photoisomerization rate (*R_Iso_*, in [*P ^∗^/*cone*/*s]) from *^P^Phi*:

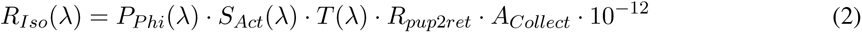

where *S_Act_* represents the activation of photoreceptors with the LEDs considering the mismatch between photoreceptor spectral sensitivity and LED emission spectrum, *T* represents the transmission of light through the mouse eye, *R*_pup2ret_ corresponds to the ratio between pupil area and retinal surface area, and *A*_Collect_ (in [*µ*m ^2^]) corresponds to light collection area of photoreceptor outer segments (for details, see Franke et al., 2019).

**Table 1:** Equipment for spatial calibration of the dome setup and building the eye tracking system.

| Part # | Name | Vendor | Part number |
| --- | --- | --- | --- |
| 1 | Laser pointer | Amazon | NC5360-944 |
| 2 | Rotation platform 1 | Thorlabs | RP005/M |
| 3 | Rotation platform 2 | Thorlabs | FP90/M |
| 4 | Rails | Thorlabs | RLA300/M and RLA450/M |
| 5 | Carrier | Thorlabs | RC2/M |
| 6 | Perpendicular carrier | Thorlabs | RC3 |
| 7 | RPI Camera | Mouser Electronics | 485-1937 |
| 8 | IR mirror | Qoptiq | G380227033 |
| 9 | IR LED | Osram | SFH 4726S |
| 10 | Raspberry Pi Version 3 Model B | Conrad Electronic | 2634243 - 62 |
| 11 | Power meter | Newport | 842-PE |
| 12 | 3D Printer | Ultimaker | Ultimaker 3 Extended |

### Eye tracking system

To investigate how mice move their eyes in response to wide-field UV and green stimuli, we recorded one or both eyes of head-fixed mice wearing a head-mounted camera system (**Fig. 1b, right**). The head-mounted eye tracking system was built based on a previous design by Meyer et al. (2018), and consisted of a Raspberry PI camera, an infrared (IR) LED, and an IR mirror. Eye tracking movies were recorded with a resolution of 640 × 480 pixels and a frame rate of 90 Hz onto the Raspberry Pi. For tracking both eyes, a second Raspberry PI camera, IR LED, and IR mirror were glued to the cap for the camera holder.

To capture eye position and pupil size in darkness under IR illumination, we first removed the IR filter from the sensor of the Raspberry PI camera (**Table 1**, part #7). The camera was then glued into a 3D printed frame, which was designed and rendered with Blender (https://www.blender.org/), prepared using Cura (Ultimaker; https://ultimaker.com/), and printed with polylactic acid (PLA) on an Ultimaker 3 Extended 3D printer. The frame consisted of a back plate covering ∼ 2*/*3 of the length of the camera and two side clamps for sliding the camera into the frame. On the top left edge of the frame, a small cube (2 × 2 mm) with a hole was used to attach a needle holding the infrared mirror (**Fig. 1b, right**).

The infrared mirror (NIR-blocking filter CALFLEX X SP, Qioptiq, UK; **Table 1**, part #8) was cut with a glass cutter into small pieces, with edge lengths of around 5–7 mm. After gluing the piece of mirror onto a small (1 × 4 mm) 3D printed tube, it was mounted on a syringe needle (Gauge 21). The needle was then bend 90° in the middle, and its tip was inserted into the hole of the cube attached to the camera frame. During implantation of the camera holder, the mirror could be moved along the needle and rotated around it, to ensure optimal positioning. The final position of the mirror was determined in a session under head-fixation, which took place after recovery from surgery and habituation to the setup and head-fixation (2 weeks post surgery). After ensuring that the camera still captured a clear and centered image of the eye, the mirror was fixed with superglue to the needle. Also in this session, the focus of the camera was adjusted, by turning the lens in its thread, before gluing it in its optimal position.

To illuminate the eye, an IR LED (Osram, SFH 4726S, Mouser Electronics, USA; **Table 1**, part #9) was glued to the bottom of the right clamp of the frame around the camera. The LED was angled towards the camera and downwards, to avoid direct reflection of the light on the mirror into the camera. The LED was powered by a Raspberry Pi single-board computer (Raspberry Pi 3 model B, Raspberry Pi Foundation, UK; **Table 1**, part #10).

The entire eye-tracking camera system was glued on an aluminum cap which tightly fitted the implanted camera holder, and whose non-symmetrical pentagon shape allowed only one particular orientation of mounting the system on the mouse’s head (**Fig. 1b, left**). The cap was fixed to the implanted camera holder by sliding a groove over the screw at the posterior side and clamping the cap by the screw-head. Clamping the two aluminum pieces with the screw secured them in a stable position over recording sessions.

### Visual stimulation

Visual stimuli were generated using custom written software in Matlab (Mathworks) within the Psychophysics Toolbox framework (Brainard, 1997; Kleiner et al., 2007). The timing of individual monitor frames was controlled by a DATAPixx input/output (I/O) hub (DATAPixx2, VPixx Technologies, Inc., Canada). This I/O hub also acquired the additional data streams and/or their timing, i.e. an analogue input representing the speed of the linear treadmill, and a TTL input signaling the acquisition time of each eye tracking camera frame. This TTL signal was sent by the Raspberry Pi that acquired the eye tracking camera frames, using the Python package `rpicamera` (Meyer et al., 2018).

#### Spatial undistortion

For spatial undistortion of the visual stimuli projected into the dome, we determined a transformation matrix, commonly referred to as the warping mesh. The transformation matrix specifies the relationship between any point on the 2D flat stimulus and any point on the dome, given the particular projector, mirror, and dome arrangement. Obtaining such a transformation matrix requires a calibration step which is known as “mesh-mapping” (Bourke, 2005; Lopes et al., 2021; Shapcott et al., 2022).

To determine the warping mesh, we displayed a polar grid whose pole was positioned at the far center of the dome, in front of the mouse. The polar grid consisted of movable points at the intersection of the concentric circles radiating out from the pole and the spokes of the grid. User interaction with these points of the grid was provided by a modified version of the function `DisplayUndistortionBVL.m` (Brainard, 1997; Kleiner et al., 2007). To span 240° in the dome, we used a grid with 8 circles and 24 spokes, with a dot spacing of 15°.

To determine the expected coordinates of the polar grid on the dome surface, we developed a tool to precisely and reversibly indicate polar angles from the center of the sphere, i.e. the position of the mouse’s head. This tool consisted of a laser pointer (**Table 1**, part #1) mounted in the center of the dome on three rotatable platforms (**Table 1**, parts #2 and #3), such that it could be rotated around three Cartesian axes. Additionally, the device was attached to a rail system (**Table 1**, parts #4–#6), which made it possible to keep the vertical and horizontal position of the laser pointer constant despite the change in angles. We used a pendulum of 60 cm length, i.e., the radius of the dome, hanging from the zenith, to re-calibrate the vertical and horizontal position of the laser pointer after rotations. One dot at a time, we positioned the laser pointer to indicate the expected angular position on the surface of the dome, and then dragged the corresponding point of the polar grid to its correct location as indicated by the laser spot on dome surface. The original and corrected pixel coordinates of the calibration points, and information about which dots were moved, were saved in a `.mat` file. This calibration file was subsequently called by a built-in function of Psychtoolbox, `CreateDisplayWarp.m`. The routine `GeometryCorrection` then applied the interpolated warping mesh to stimulus textures, thus producing stimuli with correct appearance.

### Experimental Design

To measure the pupillary light reflex (PLR; **Fig. 2**), a full-screen uniform stimulus, either UV or green, was presented in 11 linearly spaced steps ranging from 0 to 100% intensity. Each trial consisted of one intensity presented for 30 s, which was repeated 3 to 5 times in pseudo-random order.

**Figure 2:**
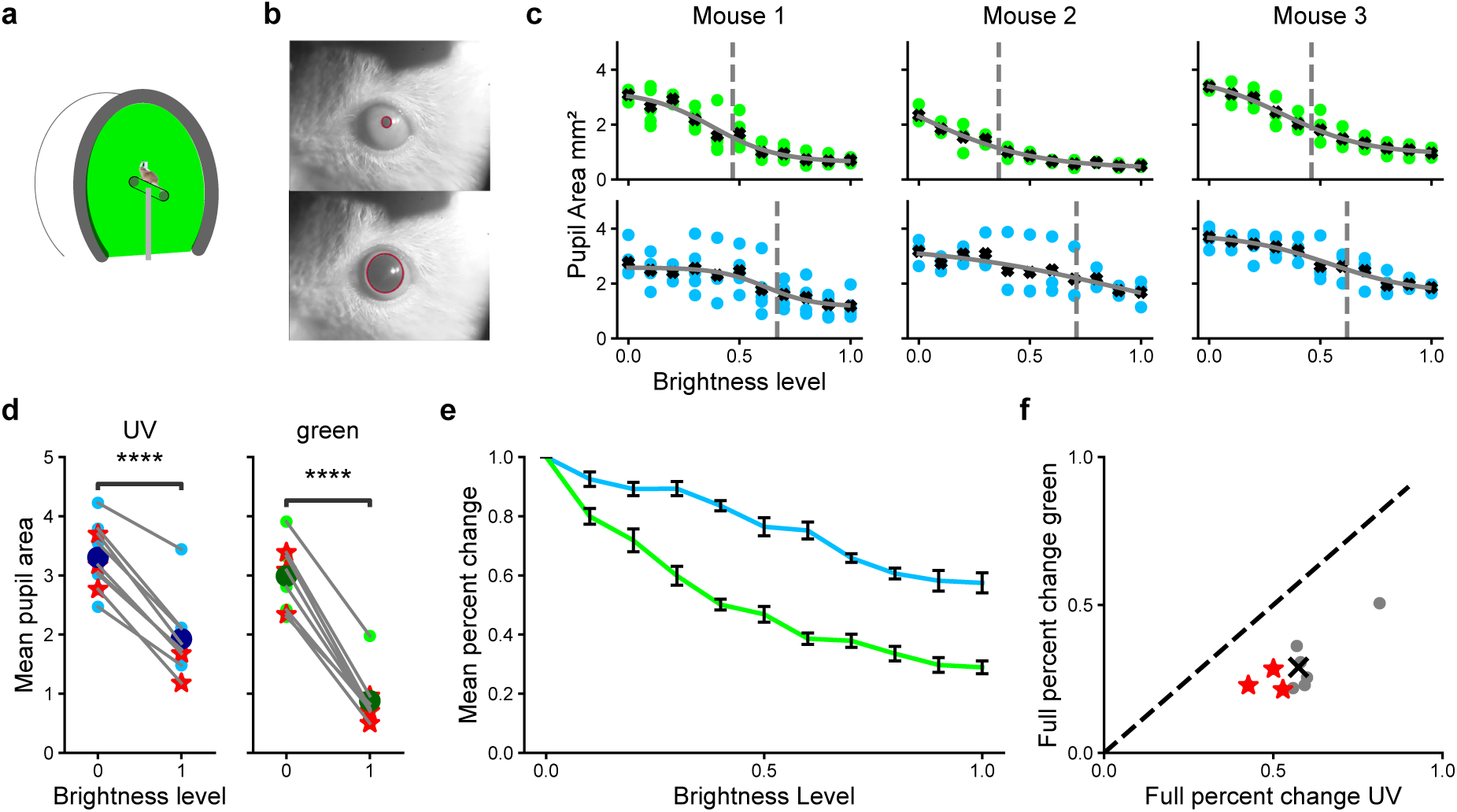
Confirmation of sufficient light intensity based on the pupillary light reflex. (**a**) Schematic drawing of the experiment: Pupil size was measured with a head-mounted eye tracking system in head-fixed mice in response to full-screen UV or green stimuli at different intensities. (**b**) Two representative video frames with fitted pupil outline (*red*) in response to 100% (*top*) and 0% (*bottom*) green light. (**c**) Pupil area as a function of light intensity for green (*top*) and UV (*bottom*); 3 mice, 2 experiments each. *Colored dots*: single trials, *black crosses*: mean across trials; *gray line*: sigmoidal fit; *gray dashed line*: light intensity where pupil area decayed to 63% of the fully dilated value. (**d**) Trial-averaged pupil area in response to zero (i.e., black screen) and full light intensity for UV (*left*) and green (*right*). *Dark blue, dark green*: Overall mean. *Red asterisks*: Example experiments shown in (c). *N* = 9 experiments in 6 mice. (**e**) Normalized change in pupil size for each light intensity for UV and green. *Error bars*: 95% CI across experiments. (**f**) Normalized change of mean pupil area between 0 and full intensity for UV and green. The dark cross represents the mean change of all experiments. *Red asterisks*: Example experiments shown in (c).

**Figure 3:**
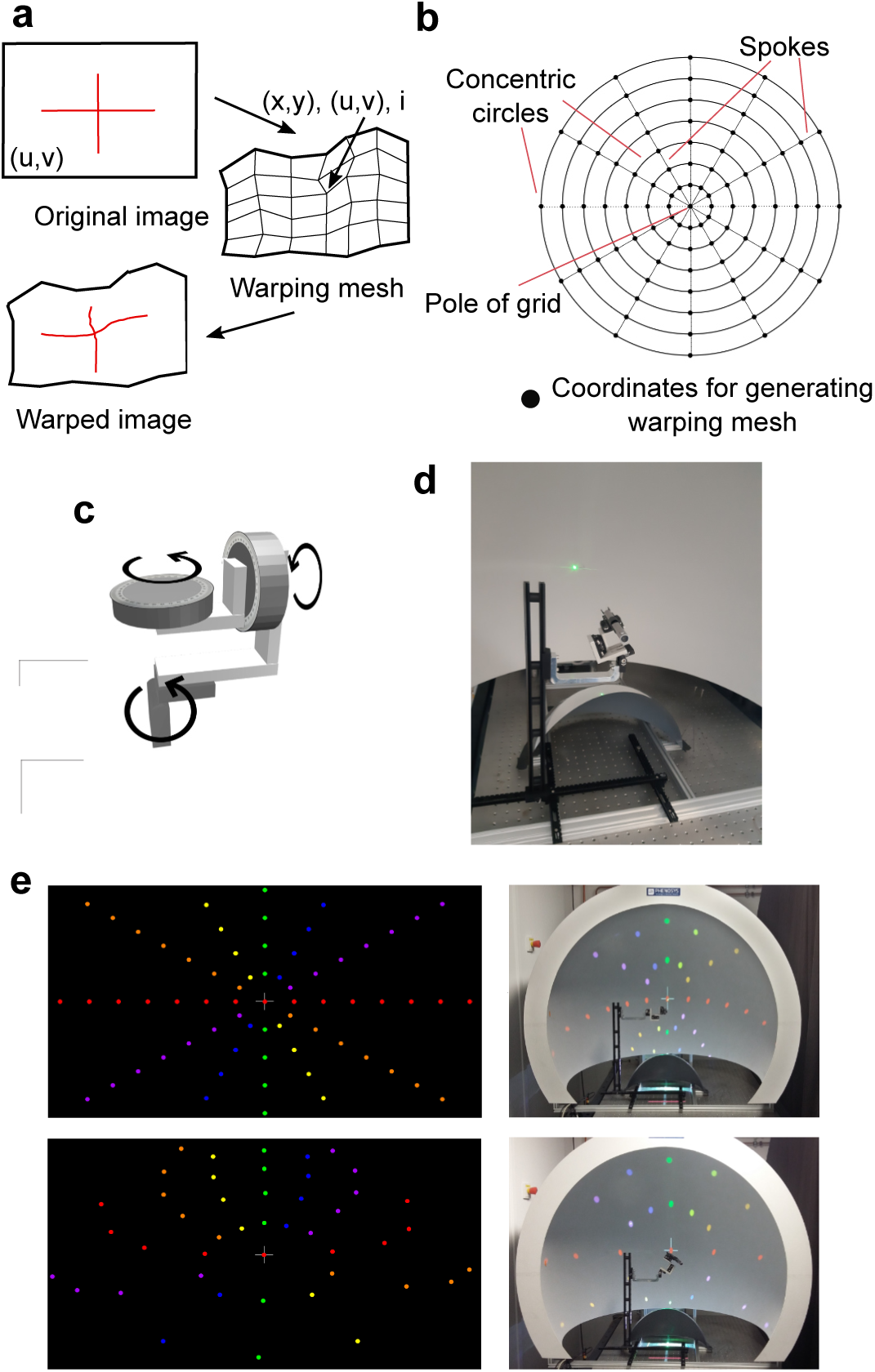
Spatial calibration of the hemispheric dome setup. (**a**) Principle of “mesh-mapping”: an image texture with coordinates (*u, v*) (*top*) is applied to a warping mesh (*right*), where each node is defined by a position (*x, y*) and texture coordinate (*u, v*). The warping (*bottom*) ensures an undistorted appearance on an arbitrary surface (adapted from: http://paulbourke.net/dataformats/meshwarp/). (**b**) Polar grid used for calibration. The pole of the grid was projected to the front center of the dome, and dots at the intersection of the concentric circles and spokes radiating from the pole were used as coordinates for generating the warping mesh. (**c**) Calibration device with 3 axis of rotation. (**d**) Calibration device mounted on rails to change its position in the sphere, allowing to keep the laser pointer tip in the center of the dome during rotations. (**e**) Polar calibration grid prior to calibration (i.e. without “mesh-mapping”) rendered on a flat screen (*top, left*) and in the dome (*top, right*). Polar calibration grid after “mesh-mapping” rendered on a flat screen (*bottom, left*) and in the dome (*bottom, right*); for illustration purposes only, the grid contained colored dots and had a 30° spacing.

For eliciting the optokinetic reflex (OKR; **Fig. 4**), a full screen drifting square-wave grating was presented. Six different gratings were presented in 90° (vertical) orientation, drifting in either clockwise or counter-clockwise directions. The drifting gratings had combinations of spatial and temporal frequencies resulting in the same speed (0.0625 cyc/° and 0.75 cyc/s; 0.125 cyc/° and 1.5 cyc/s; or 0.25 cyc/° and 3.0 cyc/s). Gratings were shown in a pseudo-random order for 1 s with an inter-trial interval of 1.5 s. Each of the 12 different conditions were repeated 50 times.

**Figure 4:**
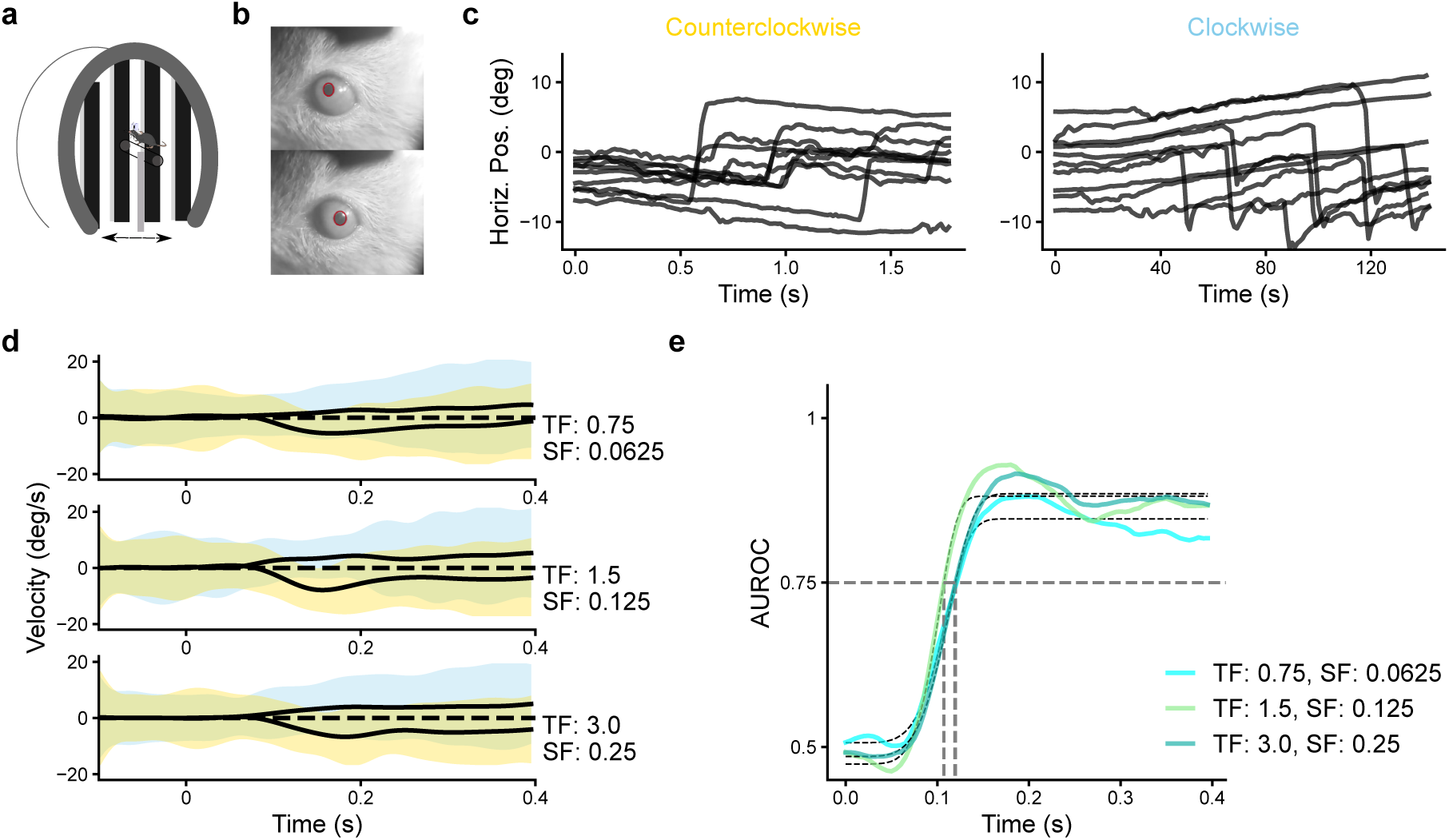
OKR measurements in the dome as a principled test for evoking eye movements in head-fixed mice. (**a**) Schematic illustration of the experiment. (**b**) Two representative video frames captured by the head-mounted eye tracking system. *Red*: fitted pupil outline. (**c**) Eye position relative to stimulus onset in one example condition (temporal frequency, TF 1.5 cyc/s, spatial frequency, SF: 0.125 cyc/deg). *Left*: counterclockwise grating drift; *right*: clockwise grating drift. (**d**) Eye velocity for all trials in all experiments (n = 7,772 trials from 26 experiments in 6 mice), separately for trials with clockwise (*blue*, standard deviation) and counterclockwise drift (*yellow*). *Black trace*: trial-averaged means. (**e**) Timepoint-by-timepoint AUROC values for the three stimulus conditions (*colored lines*) with Weibull fit (*black dashed lines*) for the extraction of latency. *Gray line*: AUROC = 0.75.

Natural movies were taken from our published open-source dataset (Qiu et al., 2021). Briefly, movies (180° FOV) were acquired, with a hand-held UV- and green-sensitive camera mounted on a gimbal, at a resolution of 437 × 437 pixels and with a frame rate of 25 frames/s. When moving the camera during the recordings, an azimuth angle of ∼ 60° between the optical axis of the fisheye lens and the movement direction was maintained, mimicking the general position of the mouse eye (Stabio et al., 2018; Oommen and Stahl, 2008). In addition, the camera was held such that the horizon bisected the camera’s visual field in elevation.

The sensitivity curve of the green and UV camera channels were determined using calibrated LEDs (green: *λ_P eak_* = 490; 525 nm, UV: *λ_P eak_* = 360; 380; 400 nm). The relationship between LED intensity and camera measurement was used to correct the intensity of the acquired movies (see Qiu et al., 2021, for details).

From this dataset (Qiu et al., 2021), we randomly selected 166 movie snippets of 10 s duration, which had been acquired during daytime and which had a mean pixel intensity across colors between 30 and 100 (out of a maximal intensity of 255). In experiments assessing pupil size during naturalistic stimulation, these 166 movie snippets were presented in two types of blocks (**Fig. 5a**). The blocks contained either the same 6 movie snippets, repeated twice per block (fixed block) or 20 random movie snippets (random block). These two types of blocks occurred in an alternating fashion, with five fixed blocks and four random blocks in one experiment. Before and after the movie sequence, the dome was illuminated for 10 min with the mean pixel intensity for UV (57) and green (61) of all individual movie snippets (blank block). In four experiments, half of the movie snippets in the fixed blocks were shown upside down, to create an unnatural distribution of chromatic input. Mice typically performed two sessions per day. The movies were presented in the dome centered at 60° azimuth, i.e. mimicking the angle during acquisition, and with a frame rate of 60 Hz.

**Figure 5:**
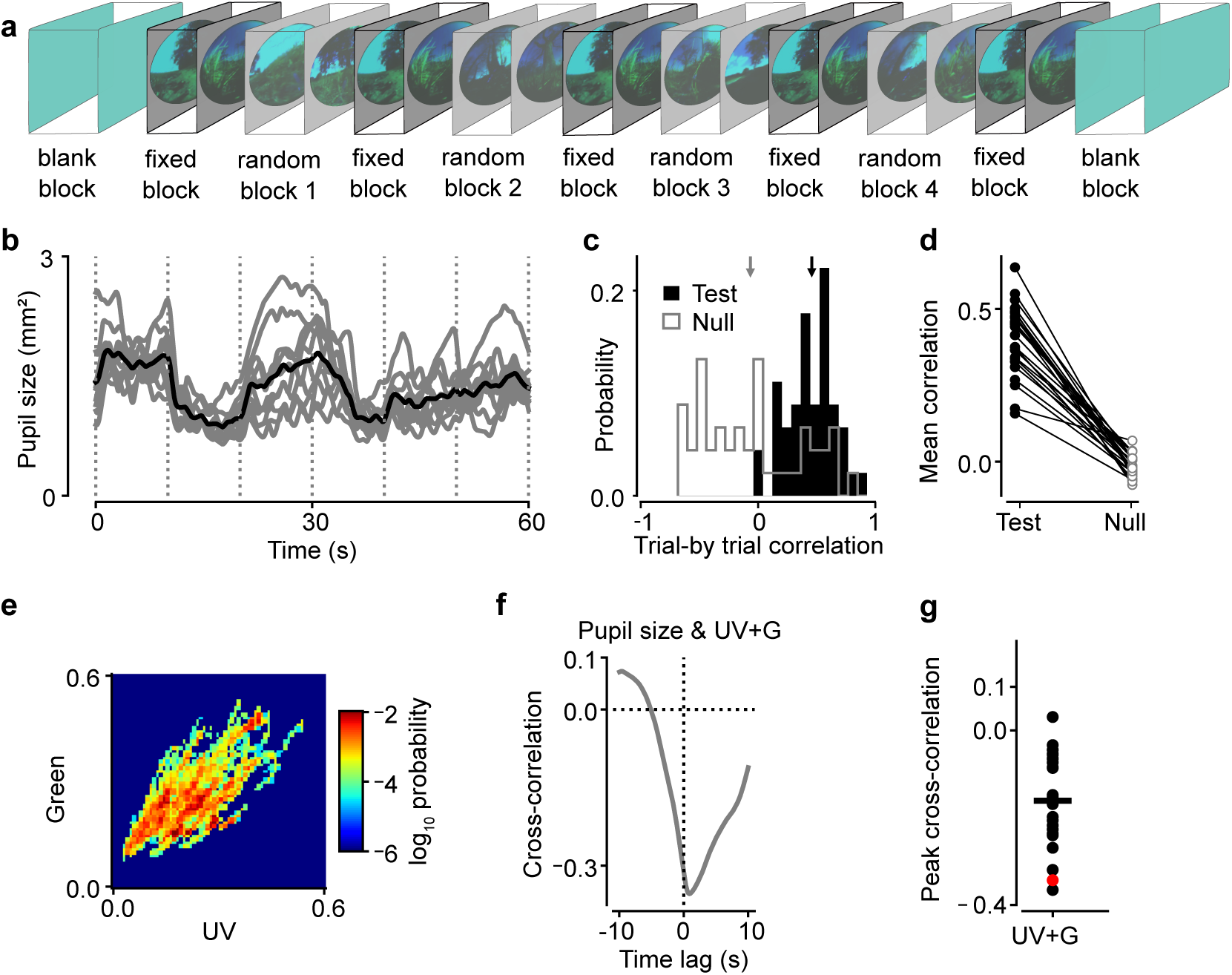
During viewing naturalistic UV/green movies, pupil size tracks average intensity of the movie frames. (**a**) Schematics of an experiment. 4 blocks of 20 random movie snippets (random block) were interleaved with a block of 6 repeated movie snippets (fixed block; snippets lasted 10 sec each). Movies were flanked by a 10 min period of blank screen of average luminance (blank block). (**b**) *Gray*: Time course of pupil size during the fixed block (example experiment, *N* = 10 repeats). *Black*: mean across repeats; *dashed vertical lines*: transitions between 10 s snippets. (**c**) Distribution of Pearson’s correlation coefficients *r* for pairs of pupil size traces in (b) (test; *black*) and for randomly selected 60 s snippets from random blocks (null; *gray*). *Arrows*: means of distributions. Same example as in (b). (**d**) Mean test and null correlations (*N* = 24 experiments from 3 mice). (**e**) 2D histogram of mean UV and green intensities for all movie frames. (**f**) Cross-correlation between pupil size and mean intensity of movie frames for both color channels combined, example experiment. Correlation at a 0.4 s lag between luminance and pupil size (Grozdanic et al., 2003; Lucas et al., 2001): -0.27 for UV + green combined. (**g**) Correlation between pupil size and mean frame intensity at a lag of 0.4 s across all experiments (*dots*). *Horizontal line*: mean, *red*: examples shown in (f).

To compare saccades and eye position between naturalistic stimulus viewing and blank blocks, we focused on experiments, in which the blank block occurred in the middle and at the end of the movie sequences (**Fig. 6b**). To minimize differential effects of behavioral adaptation, we concentrated for most of the analyses on ∼ 5 min movie blocks directly preceding the first 5 min of the blank blocks. Given the sensitivity of non-reflex behavior to numerous non-visual influences, we only reported results as significant if they occurred also in the comparison between the last 5 min of the middle blank block and the ∼ 5 min movie blocks following at position 7 and 8 in the sequence.

**Figure 6:**
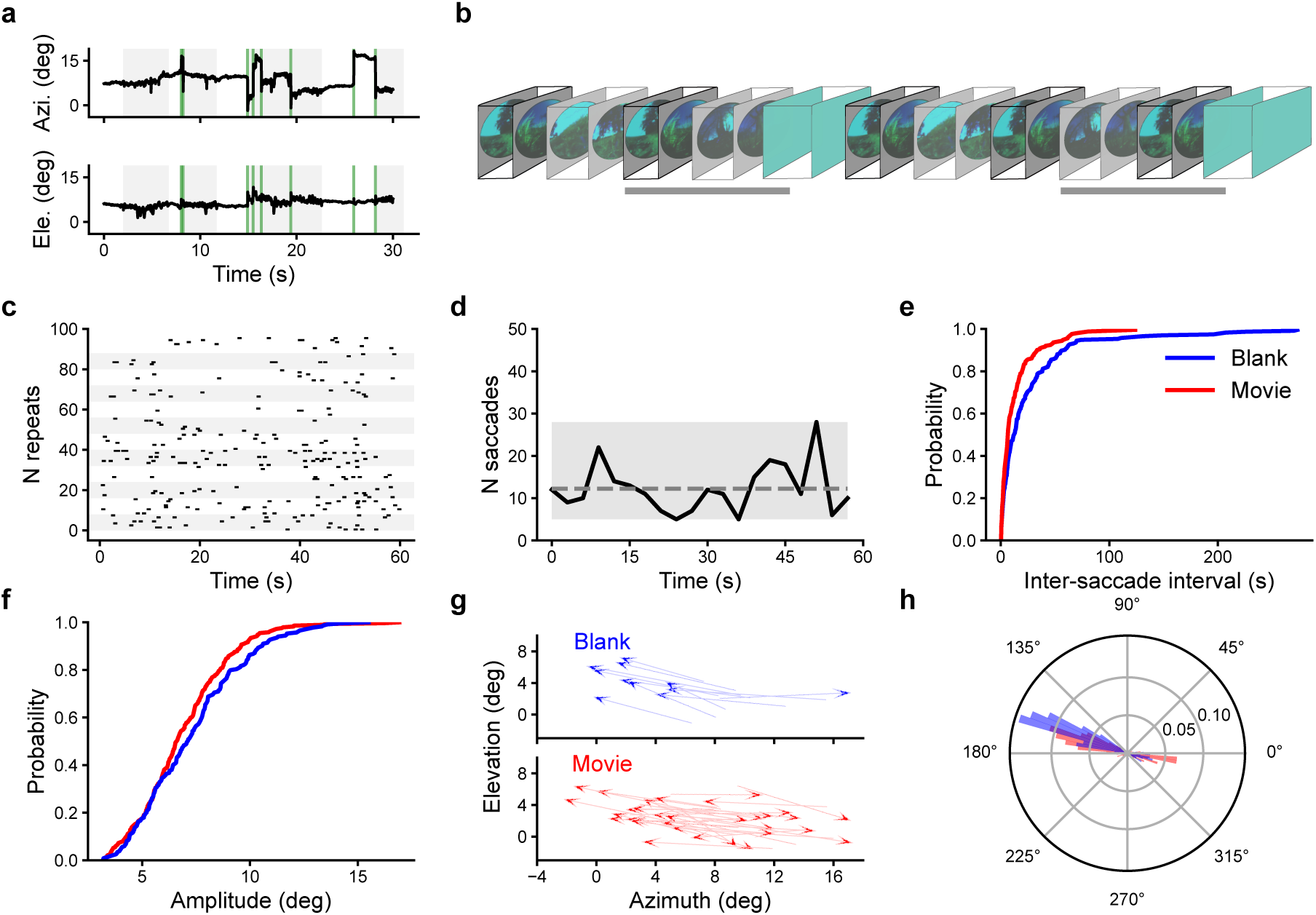
During the movie, saccadic eye movements are less bias towards nasal. (**a**) Example pupil position trace for azimuth and elevation during a movie block. *Green lines*: detected saccades; *Shaded area*: periods with running. (**b**) Schematics of an experiment, in which the blank block was presented in the middle of the movie sequence. *Horizontal gray bar*: Sequences selected for analysis. (**c**) Saccade onset times during the fixed block (*N* = 96 repeats in 3 mice). (**d**) Number of saccades across time (3 s bins), for data shown in (c). *Dashed line*: Average number of saccades per bin. *Shaded area*: 95% CI. (**e**) Cumulative distribution function of the inter-saccade intervals. (**f**) Cumulative distribution function of saccade amplitude. (**g**) Saccade vectors in an example experiment. Arrow heads indicate saccade end points. (**h**) Relative frequency of saccade directions. In (d-h), *red* refers to data from movies, and *blue* from blank blocks. (d-f, h): *N* = 235 saccades for blank blocks, *N* = 290 saccades for movies, from *N* = 12 experiments in 3 mice.

### Data Analysis

Pupil position and size, eye corners, and LED reflection position were extracted from each frame acquired by the eye tracking camera using the open-source software package DeepLabCut (DLC, Mathis et al., 2018). To train DLC to track the eye and LED markers for all experiments of all animals, we manually labeled 56 − 142 frames per mouse. Labeling entailed placing markers on the nasal and temporal eye corner, and on the edge of the pupil (8 pupil markers in total). DLC then assigned a likelihood value to each marker in each frame.

For all further analyses of the DLC output, custom code written in Python 3.8 was used. The markers around the pupil were fitted on each frame with a 2D ellipse using the scikit image class `EllipseModel`. The fit yielded parameters *cx*, *cy*, i.e. the coordinates of the ellipse center, *a*, *b*, i.e. the length of the major and minor axis, and *θ*, i.e. the orientation of the ellipse. We calculated pupil area in pixels as *area* = *πab*.

For each experiment, we first determined an appropriate threshold for the exclusion of outlier frames. We defined an outlier frame by one or more markers having a likelihood as returned by DLC of *<* 0.1 (likelihood threshold). We then randomly sampled a subset of 50 of such outlier frames per video (i.e. experiment), and visually checked the goodness of fit of the ellipse on these frames. We successively lowered the likelihood threshold until fewer than ∼ 90% of the “outlier” frames still had high quality ellipse fits. This conservative procedure led to likelihood thresholds between 0.001 and 0.1. We then used this final threshold to exclude frames with one or more pupil markers whose likelihood fell below the final threshold and linearly interpolated the fitted values of the ellipse. We additionally removed the ±10 frames adjacent to such exclusion events.

To determine pupil position, we expressed the coordinates of the ellipse center relative to the midpoint between the eye corners (with coordinate origin equaling the eye center; Meyer et al., 2020), and rotated the pixel coordinate system, such that the line connecting nasal and temporal eye corners was horizontal. We then converted pixel displacements of the pupil from the eye center into angular displacements. For the right eye, we defined positive along the horizontal axis to point away from the nose (temporal), and positive along the vertical axis to point upwards (dorsal).

To extract pupil area, we first determined the median distance between the two eye corners and assumed it to equal 3.25 mm, i.e., the default distance between the eye corners of the mouse (Schmucker and Schaeffel, 2004; Wisard et al., 2010). Using this conversion from pixels to millimeters, we could estimate the pupil area in mm^2^.

For the analysis of pupil size during measurements of the PLR, the mean pupil area per color condition was fit with a logistic function

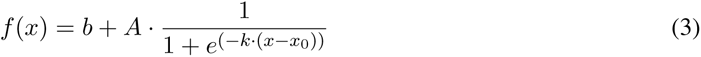

where the *b* is the baseline, *A* the overall amplitude, *x*_0_ the center, and *k* the slope of the function.

For the analysis of the OKR, we followed methods originally proposed by Tabata et al. (2011). Specifically, for all trials, the horizontal velocity of the pupil was aligned to grating drift onset. First, we upsampled the velocity traces to a frequency of 200 Hz. After excluding saccadic eye movements (acceleration *>* 200°*/*s^2^) during the first 400 ms, the velocity traces were smoothed with a digital Butterworth filter (3^rd^ order, sampling frequency = 90 Hz, low-pass, cut-off frequency = 5 Hz). To determine the latency of the OKR, for each trial, we computed eye velocities in windows of 25 ms and used a receiver operating characteristic

(ROC) analysis to quantify the separation between distributions of eye velocities for counterclockwise vs. clockwise visual motion (Tabata et al., 2011). For each time-point, AUROC (area under the ROC curve) values were computed and fit with a cumulative Weibull distribution:

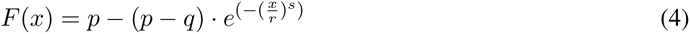

where *p* is the asymptotic level, *q* the baseline, *s* the shape, and *r* the scale parameter of the Weibull function. The latency of the OKR was then defined as the time corresponding to an AUROC value of 0.75 (Tabata et al., 2011).

For assessing the relationship between pupil size and frame intensity for natural movies, we calculated the mean intensity of each natural image frame by focusing on the pixels with y-coordinates > −35° relative to the horizon, because lower elevations were not displayed on the dome surface. We used the Pearson’s correlation coefficient to determine the relationship between mean intensity across color channels of each frame and pupil size.

To analyze rapid eye position dynamics, we extracted saccades, which we defined as eye movements with absolute velocities (first derivative) *>* 140°*/*s and amplitudes *>* 3°. To avoid double counting of saccades, we did not consider threshold crossings within the subsequent 120 ms as individual saccades. We defined the initial point and end point of each saccade as the first points, where the absolute velocity exceeded and dropped below 25°/s, respectively (Zahler et al., 2021). To analyze the consistency of saccade onset times, we divided the repeated movie sequence of the fixed block into 3 s bins and calculated the sum of saccades in each bin. In addition, we computed the average number of saccades in each bin and created 95% confidence intervals (CIs) by permuting these bins across time and repeats for 1,000 times. To assess potential difference in inter-saccade intervals (ISIs) between movies and blank blocks, we concentrated on ISIs between saccades within either movie or blank blocks. To compare distributions in ISIs and saccade amplitudes obtained during movie viewing and blank conditions, we computed a two sample Kolmogorov-Smirnov test. In addition, we calculated the medians and 95% CIs by sampling with replacement with 1,000 repeats. To test the differences in the proportions of the saccade direction, we used a Chi-square contingency test.

For the analysis of eye position during movie experiments, we subtracted from each position the mean eye position across the experiment. For all statistical tests related to eye positions, we performed two-sided, one sample permutation tests of the difference between the movie and blank conditions against 0 with 10,000 repeats.

## Results

### Hemispheric dome system with chromatic stimulation

To cover the majority of the visual field of the mouse and provide chromatic visual stimulation in the relevant spectral bands, we built a hemispheric dome setup allowing us to present wide-field stimuli in ecologically relevant “colors” (**Fig. 1**). In this setup, the mouse is head-fixed on a linear treadmill, such that its nose points to the front pole of the hemispheric screen and its head is positioned in the center of the sphere (**Fig. 1a**). A projector is placed behind and slightly below the screen and projects onto a quarter-sphere acrylic mirror positioned centrally at the bottom of the dome (**Fig. 1a,c**). Eye position of one or both eyes of the head-fixed mouse is tracked by a head-mounted eye tracking system (**Fig. 1a,b**).

As stimulus projector, we used the light guide port version of the DLP LightCrafter 4500 (LC, Texas Instruments) (**Fig. 1c,d**). Because we found that this LC model only transmits a small fraction of the light guide’s output (see also Franke et al., 2019), we took several measures to increase the light intensity available for illumination of our dome (see Methods). We chose a 6-channel high-intensity LED source (Chrolis, Thorlabs), which allowed to combine two UV LEDs (385 nm and 405 nm) and two green LEDs (475 nm and 590 nm) with different peak wavelengths for the stimulation of the UV and green mouse cone opsins, respectively (**Fig. 1e**). Translating the power of UV and green as measured from the center of the dome into photoisomerization rates, we found that our illumination provided a virtual environment at photopic conditions, with the UV LEDs activating the S-opsin up to 28, 000 *P ^∗^/*cone*/*s and the the green LEDs activating the M-opsin up to 98, 000 *P ^∗^/*cone*/*s.

To test the illumination levels of our dome using a physiological marker, we measured the pupillary light reflex (PLR, **Fig. 2**). The PLR adjusts the size of the pupil based on light intensity and thus regulates the amount of light entering the eye, with brighter light causing pupil constriction and darker light causing pupil dilation (Pennesi et al., 1998). In mice, the PLR can be driven to the same extent by both UV and green light (Yao et al., 2006). We equipped a head-fixed mouse with the head-mounted eye tracking camera, and monitored the pupil size of the right eye in response to full-screen green or UV stimuli of different intensities (**Fig. 2a**). For both green and UV stimulation, we observed pronounced decreases of pupil size with increasing light intensity (**Fig. 2b–c**). The robust negative relationship between pupil size and light intensity could be characterized by a sigmoidal curve in all mice and experiments (**Fig. 2c**). We found that the light intensity where pupil size was constricted to 63% of its fully dilated value was 36% of the maximal intensity for green and 61% for UV. Across the entire range of light intensities, the pupil constricted, on average, by 1.4 mm^2^ for UV (*p* = 1.7 × 10*^−^*^6^) and 2.1 mm^2^ for green (*p* = 2.9 × 10*^−^*^7^; **Fig. 2d**), with pupil size for the full intensity stimulus still being considerably larger for UV (1.9 mm^2^) than for green (0.9 mm^2^). Relative to a black screen, increasing light intensity successively reduced pupil size, on average, to 29% ± −2.2 (green) and 57% ± 3.4 (UV) of its initial value (**Fig. 2e,f**). Note that between wavelengths, the size of the PLR cannot be compared directly, given the ∼ 30% lower photoisomerization rate by the UV light (see previous section). Together, these results show that the illumination of the dome with both green and UV was sufficient to consistently elicit the PLR across a range of intensities.

### Spatial calibration for patterned visual stimulation

Besides achieving extensive coverage of the visual field at sufficient light intensity, a second major goal for the development of our dome setup was the ability to present arbitrary patterned visual stimuli (**Fig. 3**). In our system, displaying a flat stimulus on the curved screen of the dome is accomplished through projection onto a spherical mirror (Bourke, 2005). This mirror’s light-diverging properties enable it to reflect light over the entire dome surface. (**Fig. 1a**). However, such a projection also entails substantial, setup-specific distortions, such that a stimulus display on the dome surface with correct appearance requires a transformation of the input through a warping mesh (**Fig. 3a**).

To obtain a warping mesh specific to the distortions in our setup without prior assumptions of the underlying geometry or stimulus input, we developed an empirical calibration routine, which involved manually moving the texture coordinates (*u, v*) to the appropriate spherical coordinates (*r, θ, ρ*) (see also Lopes et al., 2021, for a similar approach). A stimulus that is well-suited for this purpose is a polar grid (**Fig. 3b**), as its correct appearance in the hemispheric dome setup is known (Bourke, 2005). Projecting the pole of the grid directly in front of the mouse, the horizontal spoke of the grid should encircle the equator of the dome, the vertical spoke should represent the meridian, and all concentric circles of the grid should be spaced regularly around the front pole in the dome (Bourke, 2005). We used a grid with 24 spokes and 8 circles, to achieve 15° of visual angles between the spokes and a coverage of 240° (**Fig. 3b and e**; for illustration purposes, only 12 spokes are shown). We displayed the intersections between the spokes and concentric circles in the polar grid as dots, which served as anchor coordinates for the calibration, because they can be easily expressed in spherical coordinates (*r, θ, ρ*) (see also Yu and Rosa, 2010) (**Fig. 3e**).

To precisely determine the location of the anchor coordinates in spherical coordinates on the dome without attaching permanent markers that might potentially interfere with the visual stimulus, we aimed a laser pointer, which was mounted on a platform allowing rotations along three axes, to specific polar coordinates on the dome’s surface (**Fig. 3c,d**). We then dragged each of the colored dots at the anchor coordinates to their expected angular position, which resulted in a set of corresponding points between the spherical coordinates (*θ, ρ*), the original image coordinates (*u, v*), and the new position of the node (*x, y*).

As expected, projecting our polar grid (**Fig. 3e**, *top left*) prior to calibration onto the dome resulted in a distorted appearance (**Fig. 3e**, *top right*). After the calibration procedure and application of the warping mesh to the grid’s texture (**Fig. 3f**, *bottom left*), the projected grid had the correct appearance in the dome (**Fig. 3f**, *bottom right*), with straight lines and regularly-spaced circles.

### Drifting gratings in the dome evoke the optokinetic reflex (OKR)

Having established methods for rendering patterned visual stimuli with correct appearance, we next sought to validate that visual stimulation in our setup can, in principle, evoke stimulus-related eye movements. We measured in head-fixed mice the optokinetic reflex (OKR), i.e., compensatory eye movements in response to large-scale retinal slip (Land, 2019).

We presented full screen, vertical gratings drifting either clockwise or counterclockwise and monitored eye position with our head-mounted, camera-based eye tracking system (**Fig. 4a**). As expected from earlier studies (Prusky et al., 2004; Stahl, 2004; Tabata et al., 2011), we observed that eye position varied systematically with the direction of drift (**Fig. 4b**). Specifically, eye movements consisted of alternating patterns of slow movements following the direction of the grating and fast, resetting movements (**Fig. 4c**). Focusing on those trials without a saccade in the first 400 ms after grating onset, we calculated eye velocity (**Fig. 4d**) and analyzed how the latency and size of the evoked eye movements depended on the temporal and spatial frequency of the drifting grating. To assess the consistency of the evoked eye motion, we pooled trials across experiments and mice (**Fig. 4d**), separated the velocities by grating drift direction, and performed an ROC analysis for each point in time (Tabata et al., 2011). The time-dependency of the resulting AUROC values was well captured by a Weibull function, and we defined the latency as the time corresponding to an AUROC value of 0.75 (**Fig. 4e**). All three stimulus conditions reliably elicited the OKR, with latencies of 115 ms ± 17 (std), 109 ms ± 15, and 118 ms ± 11, respectively (**Fig. 4e**). Together, these analyses of the OKR show that, in our dome setup, we can reliably observe stimulus-driven eye movements with the expected latency.

### Naturalistic movies elicit systematic changes in pupil size

Having established that, in principle, we can elicit and reliably track expected stimulus-evoked changes in pupil size and position, we next turned to more naturalistic visual stimulation (**Fig. 5**). More specifically, we displayed UV/green movies of outdoor scenes taken in presumed mouse habitats with a hand-held, custom camera mimicking some key aspects of the mouse visual system (Qiu et al., 2021).

An experiment consisted of several blocks of visual stimulation (**Fig. 5a**). It started and ended with a 10 min block, in which the dome was illuminated with the overall mean intensity across all individual movie frames (blank block). In between, we presented blocks containing several 10 s long movie snippets. These blocks consisted of either the same 6 movie snippets (fixed block) or 20 random snippets (random block), and occurred in an alternating fashion.

We first analyzed pupil size (**Fig. 5b**), hypothesizing that the PLR should also be elicited by fluctuations of brightness in our naturalistic movies. To validate that pupil size tracked average brightness of the movie frames, we first concentrated on the blocks containing the six repeated movie snippets. Plotting pupil size aligned to the onset of the fixed block in an example experiment indeed revealed a pattern of relatively consistent fluctuations of pupil size, occurring both between movie segments, e.g., around the transition between the first and second snippet, and within movie segments, e.g., around the middle of the fourth snippet (**Fig. 5b**). We quantified the relationship of pupil size and average brightness by correlating the time course of pupil size across pairs of trials, and found that for this experiment, the pair-wise average correlation was *r* = 0.46 ± 0.21 (mean ± std). In contrast, performing the same analysis for a randomly selected subset of 6 movie snippets from the random block yielded a correlation of *r* = −0.07 ± 0.43 (**Fig. 5c**). Repeating this analysis for all experiments (*N* = 24 experiments from 3 mice), we found an average correlation of *r* = 0.40 ± 0.12, which was significantly different from the correlation obtained with luminance input from the random movie blocks (*r* = −0.01±0.04, *p* = 0, two-sided one sample permutation test of the difference against 0; **Fig. 5d**).

Having a much greater variation of naturalistic movies beyond the fixed blocks, we next considered all movie snippets and investigated the time-course of the relationship between intensity fluctuations in UV and green, and variations in pupil size by computing the cross-correlation. Because we found UV and green intensity to be correlated in our natural movies (**Fig. 5e**) we focused on the mean intensity across spectral bands. As expected (Grozdanic et al., 2003; Lucas et al., 2001), we found a negative peak in the cross-correlation between the mean intensity of our movie frames and pupil size (−0.16 ± 0.11; **Fig. 5f,g**), indicating that the pupil constricted shortly after an increase in intensity. Taken together, our results demonstrate that fluctuations in pupil size during viewing of our naturalistic movie can be consistently related to mean frame intensity.

### Saccade direction differs for naturalistic movies and uniform screen conditions

Do head-fixed mice show differences in how they sample visual information when exposed to naturalistic movies compared to a uniform screen? We first considered saccades, i.e. rapid, conjugate eye movements which in head-fixed mice have amplitudes of 10 − 20° and peak velocities reaching 1, 000 °*/*s, and occur almost exclusively along the horizontal direction (Meyer et al., 2020; Sakatani and Isa, 2007) (**Fig. 6a**). To reduce the potential impact of factors varying over time, such as arousal or engagement with the visual stimulus, we concentrated for the subsequent analyses on experiments, in which the blank block was presented in the middle of the movie sequence and analyzed eye movements and positions during the fixed and random movie blocks immediately preceding the blank blocks (**Fig. 6b**).

To test for the absence of systematic saccades with repeated movie sequences, we first detected saccades in the fixed block. Indeed, while primates with their foveated retina explore visual scenes with predictable and repeatable saccades even during free exploration (Yarbus, 1967), saccades to target specific objects are not expected for mice or other animals with a more uniform distribution of spatial acuity (Land, 2015). Indeed, when aligning saccade times to the onset of the fixed movie block, we did not detect any systematic temporal pattern of saccade occurrence across experiments (**Fig. 6c, d**).

We next turned to the comparison of movies and blank blocks and first focused on saccade frequency. Consistent with previous reports, we found that overall saccade frequency was 0.026 saccades/s (median, 95% CI = [0.02, 0.05]; *N* = 12 experiments in 3 mice). To better understand whether these saccades occurred in specific temporal patterns that might differ between movies and blank blocks, we computed distributions of inter-saccade intervals (**Fig. 6e**). We found shorter inter-saccade intervals during movie viewing (median 6.38 s, 95% CI = [5.82, 7.65]) than blank conditions (median 11.95 s, 95% CI = [8.77, 14.58]), and hence an overall difference in their distribution (**Fig. 6e**; *p <* 0.0001, two sample Kolmogorov-Smirnov test). Given that mice exhibit a higher frequency of saccades during running (Niell and Stryker, 2010; Samonds et al., 2018), we analyzed the fraction of running, and observed that it was similar between the two conditions (median fraction running: movie 0.14, 95% CI = [0.08, 0.23], blank: 0.13, 95% CI = [0.09, 0.2]). This suggests that running is unlikely to explain the difference in inter-saccade intervals between movie viewing and blank blocks.

We then asked whether saccade amplitude and direction might differ when mice viewed naturalistic movies or a blank screen. Samonds et al. (2018) proposed that mice use saccades to refresh information: saccades during movies could be smaller and, given the dynamic stimulus input, could still direct the receptive fields of visual neurons to novel information, while saccades would need to be larger for a uniform blank input to achieve the same (see also Groner et al., 2008, for a spatial frequency effect on saccades). Contradicting this hypothesis, we found that mice made saccades of similar size during movies (**Fig. 6f**, 6.59 deg, CI = [6.38, 6.95]) compared to blanks (7.11 deg, CI = [6.66, 7.49], *p* = 0.11, two sample Kolmogorov-Smirnov test).

When considering not only amplitude, but also the direction of saccades, we found that mice had an overall bias towards nasal saccades, which was less prominent during naturalistic movie viewing than blanks. Indeed, across both stimulus classes, mice more often made saccades to nasal (62.8%) than temporal directions (23.7%; **Fig. 6g,h**). Interestingly, during movies, this bias was reduced (**Fig. 6g,h**; movie nasal 56.21% vs. temporal 28.62%; blank nasal 69.36% vs. temporal 18.72%, Chi-square test *p* = 0.004). Taken together, our analysis of saccade amplitude and direction revealed that mice make saccades of similar amplitude, but with more equal directionality during movies compared to blank periods. This pattern could reflect more active sampling of visual information during naturalistic stimulus viewing.

### Eye position is systematically shifted towards nasal for naturalistic movies

Encouraged by the observed differences in saccade direction between movies and blank conditions, we asked whether viewing of our naturalistic movies affected eye position also in general. To address this question, we again focused on the same movie and blank blocks as used above (**Fig. 6a**) and computed for each experiment distributions of eye position (**Fig. 7a,b**). Comparing eye position during naturalistic movie viewing and the blank block, we found that the average eye position during movies was more nasal (−1.12° vs. 1.19°), without substantial differences in elevation (0.08° vs. −0.08°; **Fig. 7a**). This shift towards nasal was also visible in the center of mass of the 2D distributions of eye position (**Fig. 7b**) and in their difference for movies and the blank blocks (**Fig. 7c**).

**Figure 7:**
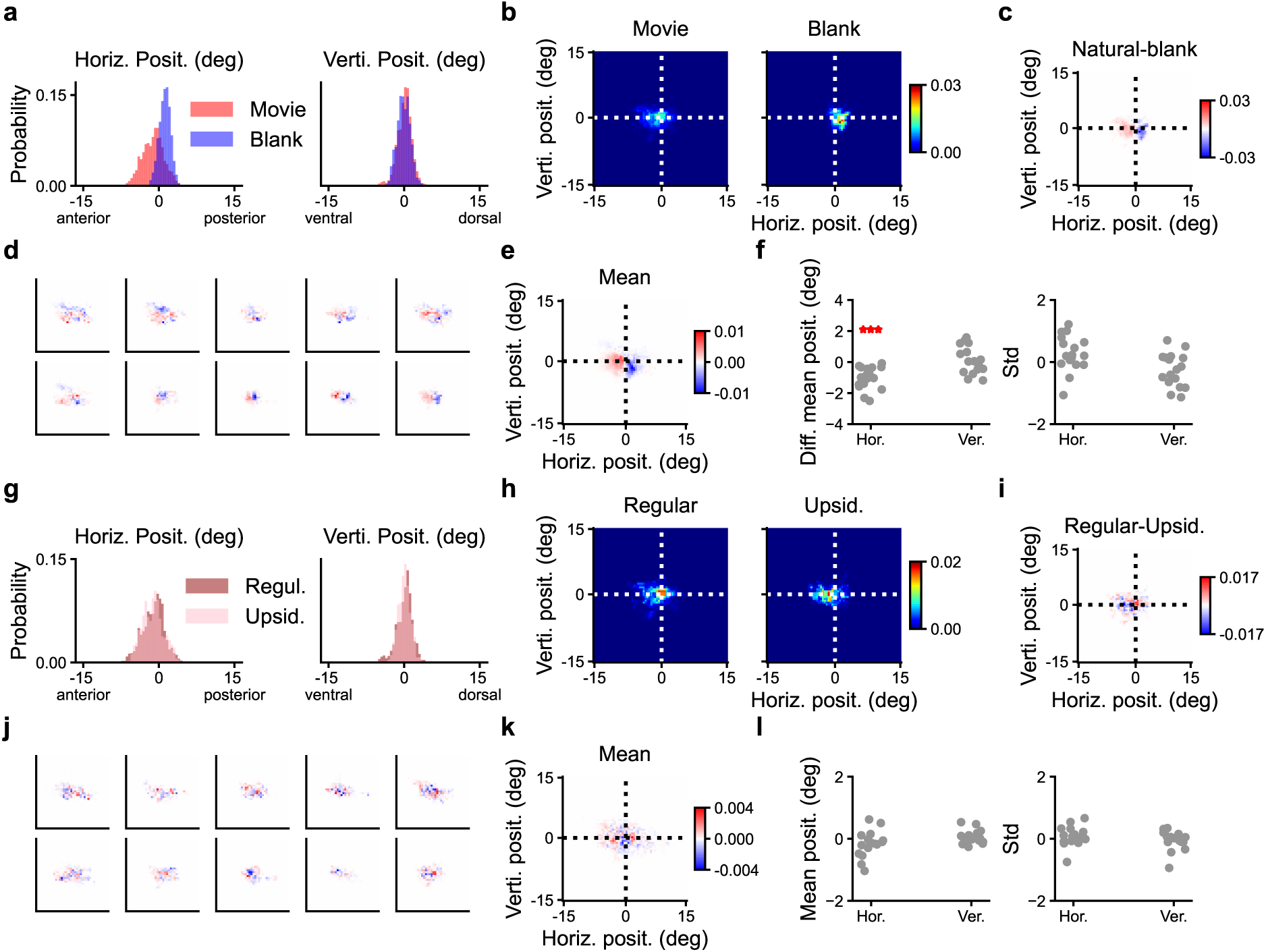
Eye position is shifted towards nasal during naturalistic movie viewing. (**a**) Distribution of horizontal (*left*) and vertical (*right*) eye position during an example experiment, separately for periods of movie viewing (*red*) and during the subsequent blank block (*blue*). (**b**) 2D probability distributions for data in (b). (**c**) Difference between 2D probability distributions of pupil position from (b) during naturalistic movie viewing vs. uniform screen (blank). (**d**) Same as (c), separately for 10 more example experiments. (**e**) Mean across all experiments. (**f**) Difference in mean (*left*) and standard deviation (*right*) of eye position distributions for naturalistic movies vs. the blank block, separately for horizontal and vertical eye position. Dots represent individual experiments. (**g**) Same as (a), separately for periods with regularly oriented movies (*red*) and upside-down movies (*pink*). (**h**) 2D probability distributions for data in (g). (**i**) Difference between the 2D distributions of pupil position from (h) during regular and upside-down movies. (**j**) Same as (i), separately for 10 more example experiments. (**k**) Mean across all experiments. (**l**) Difference in mean (*left*) and standard deviation (*right*) of eye position distributions for naturalistic movies with regular vs. upside-down orientation, separately for horizontal and vertical eye position. Dots represent the individual experiments. Data in (e-f, k-l) are from *N* = 17 experiments from 3 mice.

We found a systematic shift in average eye position not only for the example experiment, but also for individual mice and experiments (**Fig. 7d**) and in the mean across all experiments (**Fig. 7e**). Indeed, for movies compared to blank blocks, average eye position was more nasal (1.02° ± 0.72; *p <* 0.0001, two-sided one sample permutation test of the difference against 0), without concomitant differences along the vertical direction (0.10° ± 0.87, n.s.; **Fig. 7f**, left) or in the variability of eye position (movie 0.28° ± 0.57, blank −0.26° ± 0.54, n.s.; **Fig. 7f**, right). These differences in eye position did not depend on active overt behavior, as they persisted even when we restricted our analysis to periods in which the mice were quiescent (run speed *<* 2 cm/s). Together, these results indicate that during viewing of our naturalistic movies, head-fixed mice adjust their average eye position to a more nasal position.

Finally, we asked whether overall eye position of mice also changed for more subtle differences in visual input. Specifically, we hypothesized that turning the movies upside down might elicit specific changes in eye position as this manipulation reverses the vertical gradients of UV and green contrasts occurring in natural scenes (Qiu et al., 2021). To test this hypothesis, we presented the movies in half of the blocks upside-down and compared eye position between these blocks and blocks with “regular” movies. However, we did not find any systematic differences in horizontal or vertical eye position for upside-down compared to regular movies (**Fig. 7g-l**). Taken together, mice adjusted their eye position to more nasal positions when viewing our naturalistic movies compared to blank screens, but did not respond with eye position changes to more subtle manipulations of the movies.

## Discussion

Here, we describe the implementation of a hemispheric dome setup, which enables the presentation of visual stimuli in UV and green, the spectral bands relevant for mice, and provides extensive coverage of the mouse’s field of view. We verified sufficient light intensity in both wavelengths by observing systematic decreases in pupil size with increasing light intensity, i.e., the pupillary light reflex. In addition, we demonstrate that we can use the setup to elicit and capture systematic eye movements of head-fixed mice, as assessed by the optokinetic reflex to drifting gratings. We then turned to the presentation of naturalistic movies of outdoor scenes taken with a hand-held camera from the perspective of a mouse. We found that in our setup, head-fixed mice make more balanced saccades towards nasal and temporal directions during viewing of these naturalistic movies than during a uniform stimulus. In addition, head-fixed mice systematically changed their eye position during the naturalistic movies towards more nasal positions. Together, our results demonstrate that, even under head-fixation, mice perform stimulus-specific eye movements that are indicative of active sampling. We suggest that these eye movements might reflect more active engagement with dynamic naturalistic stimuli, potentially to optimize visual processing of information ahead in the visual field.

Our dome setup was inspired by previous work (e.g., Yu and Rosa, 2010; Lopes et al., 2021; Denman et al., 2017, 2018; Sibille et al., 2022; Shapcott et al., 2022; Gupta et al., 2023), exploiting hemispheric dome screens to cover large parts of the visual field of the mouse. One challenge in such setups is achieving sufficient light intensity, particularly for UV light, as the screen, mirror, and projector are typically not optimized for these wavelengths. Through a combination of a powerful LED light source and custom modifications of the projector, we were able to achieve photopic conditions. To further increase light intensity in the future, in particular in the UV range, or to accommodate a less powerful LED light source, the acrylic mirror could be exchanged against a silver-coated mirror and the dome surface could be covered with UV-reflective paint (Denman et al., 2017; Sibille et al., 2022). In addition, the size of the screen could be reduced, to decrease loss by projection distance and scatter. While the dome setup allowed us to measure behavioral responses in head-fixed mice to various stimuli, a promising further application could entail recordings of behavior and neural data in freely moving mice, similar to what has been suggested by Madhav et al. (2022).

Our experiments comparing saccadic eye movements and eye position during naturalistic movies and blank screens add to a growing literature on properties of mouse eye movements. In head-fixed mice, spontaneous saccadic eye movements are known to be rare (Niell and Stryker, 2010; Samonds et al., 2018; Meyer et al., 2020), occur mainly in the horizontal direction (Sakatani and Isa, 2007; Meyer et al., 2020), and seem to happen mostly during attempted head-movements (Meyer et al., 2020; Zahler et al., 2021). A previous study showed that under head-fixation, directionally biased saccades and attempted head-movements can be triggered in mice by tactile stimuli (Zahler et al., 2021). Our results suggest that also different classes of visual stimuli can elicit saccades with different directionality: while we observed a dominance of nasally-directed saccades during blank screens, saccades during naturalistic movies occurred in both nasal and temporal directions in a more balanced way. This might reflect an active sampling of information during patterned visual input.

Besides visual orienting, another proposed function of saccades could be that they de-correlate the visual input (Samonds et al., 2018). Consistent with this proposal, it has been found that the amplitude and frequency of saccades in mice depend on the size or spatial frequency of the visual stimulus (Samonds et al., 2018), with larger and more frequent saccades during larger or more low spatial-frequency input. Similarly, a study in humans showed less frequent and smaller eye movements during viewing of dynamic stimuli, where information changes over time irrespective of eye movements, compared to static visual scenes (Dorr et al., 2010). Under our visual stimulation conditions, comparing between naturalistic movies and blank stimuli, we found no difference in saccade size, or in variability of general eye position. One potential explanation for this discrepancy could be that naturalistic movies and blanks, in addition to differing in spatial frequency content and dynamics, likely lead to different levels of engagement, which might override some of the stimulus-specific effects. Undeniably, a uniform screen and a dynamic naturalistic movie are at opposite ends of the spectrum in terms of several stimulus parameters. For future experiments it would thus be interesting to vary the visual stimulus in more fine-grained ways, e.g. by creating various types of noise with defined spatio-temporal spectra, in order to test some of the above interpretations and allow a more direct comparison to the previous literature.

What could be the purpose of the systematic change in eye position towards nasal directions that we observed during viewing of a naturalistic stimuli? While originally it was thought that the mouse retina contained topologically uniform functional feature selectivity, recent years have revealed several retinal specializations (reviewed in Sedigh-Sarvestani and Fitzpatrick, 2022). For instance, sustained alpha retinal ganglion cells display a nasal-to-temporal gradient in cell density, size, dendritic signal integration, and receptive fields, which might serve to enhance visual sampling of frontal space (Bleckert et al., 2014; Oesterle et al., 2024). In addition, in mouse visual cortex, neurons representing the frontal space have smaller receptive fields (van Beest et al., 2021) and encode binocular disparity combining the information from both eyes (La Chioma et al., 2019; Samonds et al., 2019; Scholl et al., 2013). Furthermore, behavioral studies in head-fixed mice have revealed higher visual acuity for this frontal space of enhanced neural coverage (van Beest et al., 2021). Finally, freely moving mice use gaze changes to move prey objects into the central portion of their visual field (Holmgren et al., 2021; Johnson et al., 2021; Michaiel et al., 2020; van Beest et al., 2021). Thus, positioning the eyes in a more frontal position during viewing of naturalistic movies might increase binocular overlap and direct higher-acuity processing resources to the field of view straight ahead.

Arguably the best studied specialization in the mouse retina is the gradient of cone opsin expression, with a relatively uniform distribution of S-cones, and a dorso-ventral gradient of S-cone opsin co-expression in the M-cones (Szél et al., 1992; Baden et al., 2013; Nadal-Nicolás et al., 2020). This arrangement results in a strongly UV-sensitive ventral and a more green-sensitive dorsal retina, which closely mirrors the relative abundance of contrasts in natural scenes (Qiu et al., 2021), and thus seems to support the detection of dark contrasts in the sky, such as aerial predators (Qiu et al., 2021; Baden et al., 2013). When artificially inverting the UV-green gradient by turning the input movies upside down, we did not observe any systematic differences in overall eye position or other behavioral response. Future experiments in freely moving mice could test whether a more subtle manipulation, e.g. an up- or downwards shift of the naturalistic movies and thus the horizon might lead to concomitant changes in gaze, potentially to align the retinal gradient of cone opsin expression to the horizon.

Taken together, our hemispheric dome setup, capable of presenting naturalistic visual stimuli with extensive coverage and in the relevant spectral bands for mice, allowed us to discover stimulus-specific patterns of eye movements and adjustments of eye position, indicative of active visual sampling in head-fixed mice.

## Author Contributions

LB, TE, MK, YQ, VP Designed Research; MK, VP Performed Research; YQ, ZZ, SK, AK Contributed unpublished reagents/ analytic tools; MK, VP, YQ, Analyzed data; LB, TE, SK, MK, VP, YQ Wrote the paper

## Acknowledgments

This research was supported by the Deutsche Forschungsgesellschaft (DFG) Sonder-forschungsbereich (SFB) 1233, *Robust Vision: Inference Principles and Neural Mechanisms*, Teilprojekt (TP) 10, project number: 276693517 (L.B., T.E.). We thank A. Meyer for help with establishing head-mounted eye tracking, D. Crombie for software for behavioral analyses, and F. Schneider-Soupiadis for support with code. We also thank T. Wachtler for advice and sharing of equipment used for calibrating the dome, and A. Genewsky for useful suggestions and discussion about hardware development. Thanks also go to M. Sotgia for lab management and support with mouse handling, S. Schörnich for IT support, and B. Grothe for providing an excellent research infrastructure.

